# *In Vitro* efficacy comparison of linezolid, tedizolid, sutezolid and delpazolid against rapid growing Mycobacteria isolated in Beijing, China

**DOI:** 10.1101/2020.06.25.172742

**Authors:** Shuan Wen, Xiaopan Gao, Weijie Zhao, Fengmin Huo, Guanglu Jiang, Lingling Dong, Liping Zhao, Fen Wang, Xia Yu, Hairong Huang

**Affiliations:** National Clinical Laboratory on Tuberculosis, Beijing Key laboratory for Drug-resistant Tuberculosis Research, Beijing Chest Hospital, Capital Medical University, Beijing, China; MOH Key Laboratory of Systems Biology of Pathogens, Institute of Pathogen Biology, Chinese Academy of Medical Sciences & Peking Union Medical College, Beijing, China; The administration office of clinical trial, Beijing Chest Hospital, Capital Medical University, Beijing, China

**Keywords:** Rapidgrowing Mycobacteria, delpazolid, sutezolid, tedizolid, linezolid, antimicrobial activity

## Abstract

The natural resistance of rapid growth Mycobacterium (RGM) against multiple antibiotics renders the treatment of caused infections less successful and time consuming. Therefore, new effective agents are urgently needed. The aim of this study was to evaluate the *in vitro* susceptibility of 115 isolates, constituting different RGM species, against four oxazolidinones i.e. delpazolid, sutezolid, tedizolid and linezolid. Additionally, 32 reference strains of different RGM species were also tested. The four oxazolidinones exhibited potent *in vitro* activity against the recruited RGM reference strains, 24 out of 32 RGM species had MICs ≤ 8 μg/mL against all four oxazolidinones whereas tedizolid and delpazolid generally presented lower MICs than linezolid or sutezolid. Tedizolid showed the strongest activity against clinical isolates of *M. abscessus* with MIC_50_=1μg/mL and MIC_90_=2μg/mL. MIC values for tedizolid were usually 4- to 8-fold less than the MICs of linezolid for *M. abscessus* subsp. *abscessus*. The MIC distributions of sutezolid and linezolid were similar, while delpazolid showed 2-fold lower MIC as compared with linezolid. Linezolid was not active against most of the tested *M. fortuitum* isolates, 22 out of the 25 *M. fortuitum* were resistant against linezolid. However, delpazolid exhibited better antimicrobial activity against these isolates with 4-fold lower MIC values, in contrast with linezolid. In addition, the protein alignment of RplC and RplD and structure based analysis showed that there may be no correlation between oxazolidinones resistance and mutations *in rplC, rplD* and *23 srRNA*genes in tested RGM. This study showed tedizolid harbors the strongest inhibitory activity against *M. abscessus in vitro*, while delpazolid presented the best activity against *M. fortuitum*, which provided important insights on the potential clinical application of oxazolidinones to treat RGM infections.

## INTRODUCTION

Non-tuberculous mycobacteria (NTM) are recognized as important opportunistic pathogens of humans that can cause pulmonary infection, lymphadenitis, skin abscesses, disseminated infection and systematic infection. The prevalence of NTM infections has increased globally and even surpassed tuberculosis (TB) in certain countries (1–5). According to their speed of growth (i.e. appearance of visible colonies within or after 7 day cultivation on solid medium), NTM can be categorized as rapid growing Mycobacteria (RGM) or slow growing mycobacteria (SGM). Compared with SGM, RGM are more resistant to conventional anti-TB agents and other general antibiotics, therefore, increasing the chances of treatment failure(6). *M.abscessus* and *M.fortuitum* are among the most frequently isolated and pathogenic RGMs(1–5). *M.abscessus* often cause severe pulmonary infections with poor clinical outcomes and have been frequently reported to cause soft tissue infections(7). *M.fortuitum* can cause soft tissue infection during trauma and surgery, while lung disease caused by them is rare(8). The limited efficacies and availability of only fewer choices of medications highlight the requirement of identifying new and more potent antimicrobials against RGMs.

Oxazolidinones have demonstrated promising efficacies against *M.tuberculosis* (TB) *in vitro* and *in vivo*. Due to their distinct mechanism of action (binding to the 23S ribosome, thereby blocking microbial protein synthesis) without cross-resistance to the existing TB drugs. Oxazolidinones are proposed to be used for the treatment of multiple drug resistant TB. Linezolid (LZD), licensed in 2000, is an oxazolidinone which exhibited excellent antibacterial activity against drug resistant tuberculosis (DR-TB) and NTM infection(9–12). However, serious hematologic and neurologic toxicities can be caused by LZD during long term therapy due to its inhibition of mitochondrial protein synthesis which often requires dose reduction or discontinuation(13). Thus, new oxazolidinones drugs with superior efficacy and reduced toxicity are continuously sought.

Recently, three new next-generation oxazolidinones have been developed for potential use against DR-TB. Tedizolid (TZD) phosphate is a novel, potent oxazolidinone pro-drug that has been approved by the American FDA (2014) and the European Medicine Agency (2015) for treatment of acute bacterial skin and soft tissue infections(7). The pharmacokinetic/pharmacodynamics properties of TZD allow it to be administered orally once daily, facilitating its usage in prolonged treatment course. Sutezolid (SZD) (PNU-100480) is a thiomorpholinyl analog of LZD with preliminary evidence of superior efficacy against *M. tuberculosis*(14). SZD was found to be generally safe, well tolerated in TB patients, and with readily detectable bactericidal activity in sputum and blood. Delpazolid (LCB01-0371)(DZD) is a thiomorpholinyl analog of LZD, which showed superior efficacy against *M.tuberculosis* in the hollow-fiber, mouse model, and whole-blood model (2–4). DZD was well tolerated and showed bacteriostatic and bactericidal activity comparable to LZD against *S. aureus*, *E. faecalis* and methicillin-resistant *Staphylococcus aureus* in a recently completed phase I clinical trial(15).

To better understand the efficacies of these three new-generation oxazolidinones against different RGM species, we detected the MICs of 32 RGM reference strains and 115 RGM clinical isolates collected in Beijing, China. Furthermore, we investigated the three reported LZD-resistance genes (including *rplC, rplD* and *23srRNA*) from different RGM species to identify their potential relationships with oxazolidinone resistance.

## RESULTS

### MICs of SZD, TZD, DZD and LZD against RGM reference strains

The MICs of the 32 reference strains against SZD, TZD, DZD and LZD are presented in Table 1. All four oxazolidinones exhibited antimicrobial activities *in vitro* against the recruited RGM reference stains. Majority of the species had MICs equal to or below 8μg/mL for all four drugs. Only *M.fortuitum* and *M.rhodesiae* had MICs greater than 32μg/mL. Generally, a given isolate presented uniform tendency against all four oxazolidinones, the MIC values were either high for or low for the four drugs. For *M.abscessus*, the efficacy of TZD was stronger than LZD.

**Table 1.** MICs of LZD, TZD, SZD and DZD against reference strains of 32 RGM species

| Strain by type | Mycobacterium species<br>(strain) | MIC(µg/ml) |  |  |  |
| --- | --- | --- | --- | --- | --- |
|  |  | LZD | TZD | SZD | DZD |
| RGM species |  |  |  |  |  |
| ATCC 19977 | *Mycobacterium abscessus* | 16 | 4 | 8 | 8 |
| ATCC 27406 | *Mycobacterium agri* | 0.25 | 0.25 | 0.25 | 0.5 |
| ATCC 27280 | *Mycobacterium aichiense* | 0.5 | 0.5 | 2 | 1 |
| ATCC 23366 | *Mycobacterium aurum* | 0.25 | 0.125 | 0.25 | 0.125 |
| ATCC 33464 | *Mycobacterium*<br>*austroafricanum* | 0.25 | 0.25 | 2 | 0.5 |
| ATCC 14472 | *Mycobacterium chelonae* | 8 | 8 | 4 | 8 |
| ATCC 19627 | *Mycobacterium chitae* | 1 | 1 | 2 | 1 |
| ATCC 27278 | *Mycobacterium chubuense* | 32 | 2 | 2 | 2 |
| DSM 44829 | *Mycobacterium cosmeticum* | 4 | 2 | 16 | 1 |
| ATCC 19340 | *Mycobacterium diernhoferi* | 1 | 0.5 | 2 | 1 |
| ATCC 43910 | *Mycobacterium duvalii* | 1 | 0.25 | 0.5 | 0.5 |
| ATCC 35219 | *Mycobacterium fallax* | 4 | 2 | 8 | 1 |
| ATCC 14474 | *Mycobacterium flavescens* | 16 | 2 | 2 | 8 |
| ATCC 6841 | *Mycobacterium fortuitum* | 32 | >32 | >32 | >32 |
| ATCC 27726 | *Mycobacterium gadium* | 0.5 | 0.125 | 0.25 | 0.125 |
| ATCC 43909 | *Mycobacterium gilvum* | 0.5 | 0.25 | 0.5 | 0.5 |
| ATCC<br>BAA-955 | *Mycobacterium goodii* | 16 | 4 | 32 | 2 |
| DSM 44124 | *Mycobacterium mucogenicum* | 1 | 1 | 1 | 1 |
| ATCC 25795 | *Mycobacterium neoaurum* | 1 | 0.5 | 2 | 1 |
| ATCC 27023 | *Mycobacterium obuense* | 0.5 | 0.25 | 0.5 | 0.5 |
| ATCC 19686 | *Mycobacterium parafortuitum* | 1 | 0.5 | 2 | 1 |
| DSM 43271 | *Mycobacterium peregrinum* | 2 | 4 | 4 | 1 |
| ATCC 11758 | *Mycobacterium phlei* | 2 | 4 | 8 | 16 |
| ATCC 33776 | *Mycobacterium porcinum* | 16 | 8 | 32 | 4 |
| ATCC 35154 | *Mycobacterium pulveris* | 1 | 1 | 1 | 2 |
| ATCC 27024 | *Mycobacterium rhodesiae* | >32 | >32 | >32 | 16 |
| ATCC 700731 | *Mycobacterium septicum* | 16 | 8 | 8 | 8 |
| ATCC 19420 | *Mycobacterium smegmatis* | 2 | 2 | 4 | 4 |
| ATCC 19527 | *Mycobacterium*<br>*thermoresistibile* | 4 | 2 | 2 | 4 |
| ATCC 27282 | *Mycobacterium tokaiense* | 1 | 2 | 2 | 1 |
| ATCC 23292 | *Mycobacterium triviale* | 2 | 2 | 2 | 2 |
| ATCC 15483 | *Mycobacterium vaccae* | 2 | 0.5 | 1 | 1 |

### The MIC distributions of M.abscessus and M.massiliense against LZD, TZD, SZD and DZD

The MIC distributions of *M.abscessus* and *M.massiliense* against LZD, TZD, SZD and DZD are shown in Figure 1. MICs for TZD were generally 4- to 8-fold less than the MICs of LZD for the two species. The MIC distribution of SZD was similar to LZD, while DZD values were generally half of LZD. Notably, TZD showed strongest activity against *M.abscessus* with MIC_50_=1μg/mL and MIC_90_=2μg/mL. According to the CLSI resistance criteria for LZD(16), the susceptibility rate of *M.abscessus* against LZD, TZD, SZD and DZD was 73.5%(36/49), 100%(49/49), 71.4%(35/49), 87.8%(43/49), respectively. The susceptibility rate of *M.massiliense* against LZD, TZD, SZD and DZD was 65.8%(23/35), 82.9%(29/35), 68.6%(24/35) and 74.3%(26/35), respectively. In general, the MIC distributions of *M.massiliense* had uniform tendency than *M.abscessus*, but with an exception for TZD. The MICs of *M.massiliense* isolates were higher than *M.abscessus*. 6 out of 35 isolates of *M.massiliense* had MICs ≥ 16μg/mL against TZD, the MICs of all the tested *M.abscessus* were ≤4μg/mL. In addition, the MIC outcomes for species with less than five isolates are presented in Table 2.

**Figure 1.**
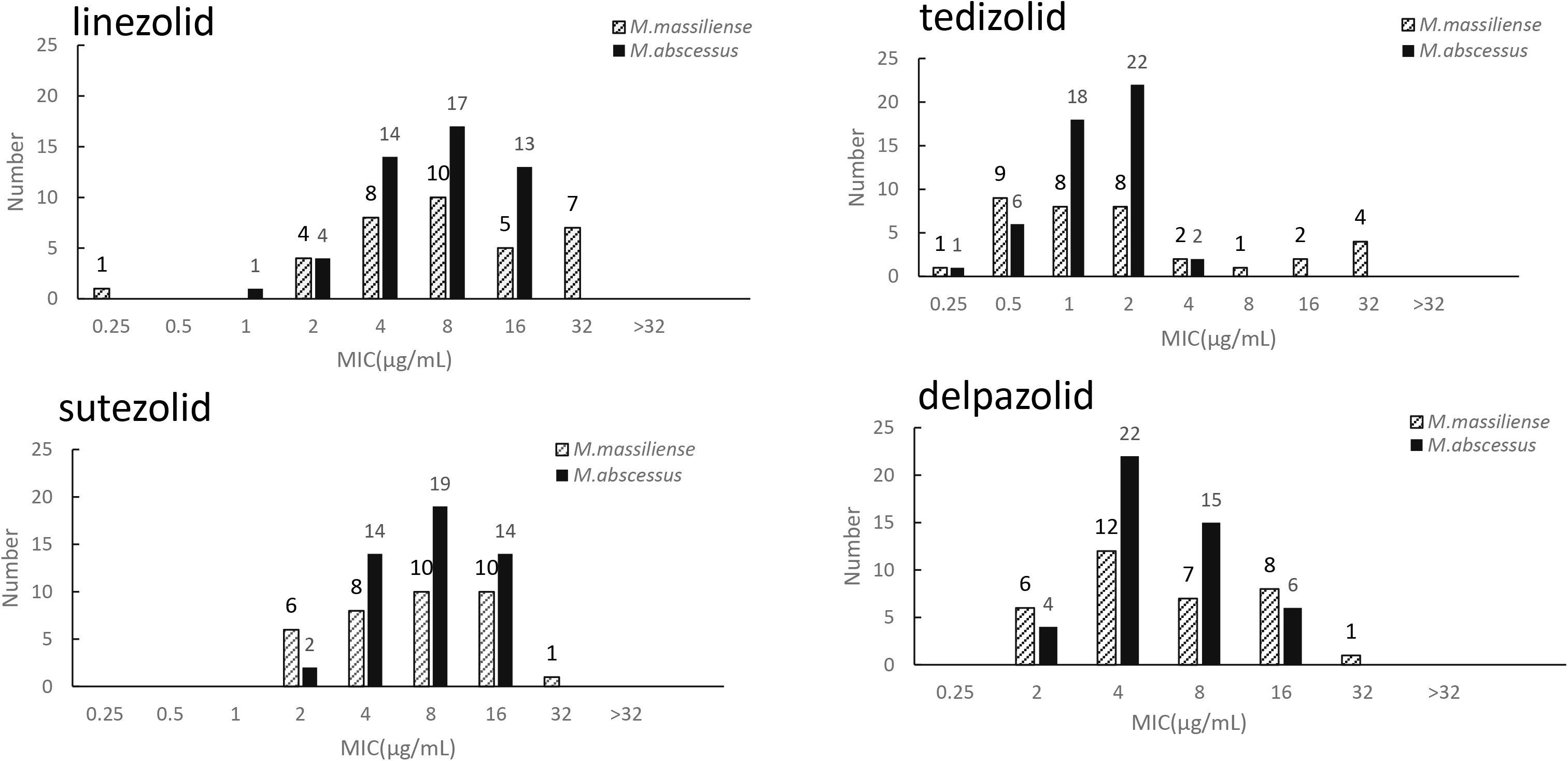
The MIC distributions of *M. abscessus* and *M. massiliense* against LZD, TZD, SZD and DZD.

**Table 2.** MICs of LZD, TZD, SZD and DZD against reference strains of 6 RGM species

| Mycobacterium species<br>(strain) | Clinical isolates<br>number | MIC(μg/ml) |  |  |  |
| --- | --- | --- | --- | --- | --- |
|  |  | LZD | TZD | SZD | DZD |
| RGM species |  |  |  |  |  |
| Mycobacterium chelonae | 585 | 32 | 8 | 16 | 8 |
| Mycobacterium chelonae | 752 | >32 | 16 | 8 | 16 |
| Mycobacterium chelonae | 1354 | 4 | 1 | 2 | 1 |
| Mycobacterium chelonae | 1392 | 4 | 1 | 2 | 1 |
| Mycobacterium chelonae | 1593 | 4 | 0.5 | 2 | 0.5 |
| Mycobacterium porcinum | 29891 | 4 | 2 | 4 | 1 |

### The MIC distributions of M.fortuitum against LZD, TZD, SZD and DZD

The MIC distributions of *M.fortuitum* against LZD, TZD, SZD and DZD are shown in Figure 2. In contrast to *M.abscessus* and *M.massiliense*, *M.fortuitum* presented higher percentage of resistance against the four oxazolidinones. The susceptibility profiles of the clinical isolates exhibited much lower MICs than *M.fortuitum* ATCC6481 reference strain. In Total, 88%(22/25) of the clinical isolates were resistant to LZD, including *M.fortuitum* reference strain. The *in vitro* activity of DZD was relatively better than LZD as indicated by its 2- to 4-fold lower MIC. The MIC distributions of TZD was similar to LZD as only 5 out of 25 isolates indicated MIC≤8μg /mL. According to the cutoff value of LZD, the susceptibility rates of *M.fortuitum* against TZD, SZD and DZD were 20%(5/25), 12%(3/25), 76%(19/25), respectively.

**Figure 2.**
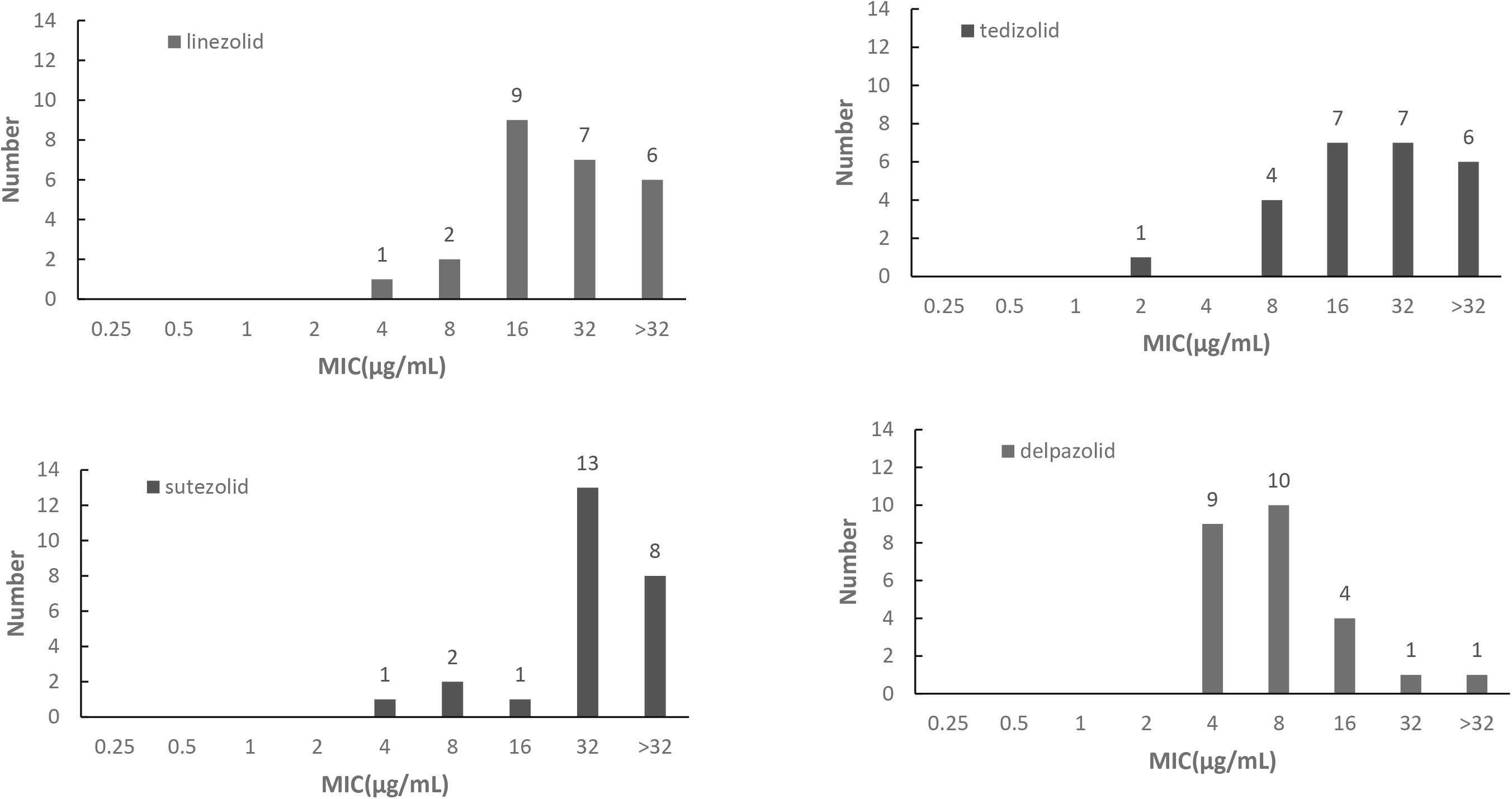
The MIC distributions of *M. fortuitum* against LZD, TZD, SZD and DZD.

### Alternations in the Oxazolidinones target sites

The entire *23SrRNA*, *rplC*, and *rplD* genes were sequenced to identify the potential mutations associated with oxazolidinones resistance. The sequences of the tested clinical isolates of *M.abscessus, M.massiliense* and *M.fortuitum* were compared with their corresponding reference strains. For *M.massiliense* isolates, Ala177Proin *rplD* was detected in 12 isolates with MIC of LZD ≥2μg/mL. In addition, two types of synonymous SNPs within the coding region of *rplC* were also observed both in LZD resistant and susceptible isolates, including Leu86Leu(CTG → CTT) and Ala92Ala(GCG→GCT). A2271G in 23SrRNA was found in one isolate with MIC of LZD=8μg/mL. For *M.abscessus* isolates, no non-synonymousmutation in the coding gene of *rplC* and *rplD* was observed, while most frequently observed mutation i.e. T2650C(n=2) was found in *23SrRNA* with MIC of LZD ≥2μg/mL(Table 3).

**Table 3.** The MICs of LZD and *rplC*, *rplD* and *23srRNA* mutations against *M. abscessus* and *M. massiliense* isolates

| MIC of LZD (μg/ml) | Species (NO.) | RplC | RplD | 23SrRNA |
| --- | --- | --- | --- | --- |
| 0.25 | *M. abscessus*(0) | — | — | — |
|  | *M. massiliense*(1) | Leu86Leu(1) | Gly75Gly(1) | WT |
| 1 | *M. abscessus*(1) | WT | WT | WT |
|  | *M. massiliense*(0) | — | — | — |
| 2 | *M. abscessus*(4) | WT | WT | G1914A (1)<br>T2650C(1) |
|  | *M. massiliense*(4) | Leu86Leu(2) | Gly75Gly(2)<br>Ala177Pro(2)<br>Val192Val(2) | WT |
| 4 | *M. abscessus*(14) | WT | Phe23Phe(2) | T2650C(1) |
|  | *M. massiliense*(8) | Leu86Leu(5) | Gly75Gly(5)<br>Ala177Pro (3)<br>Val192Val(3) | G2582C(1)<br>-2625AC(1) |
| 8 | *M. abscessus*(17) | WT | Phe23Phe(3)<br>Gly111Gly(1) | WT |
|  | *M. massiliense*(10) | Leu86Leu(4)<br>Ala92Ala(2) | Ile29Ile(1)<br>Gly75Gly(3)<br>Ala177Pro (4)<br>Val192Val(6) | A2271G(1) |
| 16 | *M. abscessus*(13) | WT | Phe23Phe(3) | WT |
|  | *M. massiliense*(5) | Leu86Leu(2) | Gly75Gly(2)<br>Ala177Pro (3)<br>Val192Val(3) | WT |
| >16 | *M. abscessus*(0) | — | — | — |
|  | *M. massiliense*(7) | Leu86Leu(1)<br>Ala92Ala(6) | Gly75Gly(1)<br>Val192Val(5) | WT |

Among the tested *M.fortuitum* isolates, all MICs for LZD were above 2μg/mL. No non-synonymous mutation was detected in the *rplC* gene. Among 25 tested *M.fortuitum* isolates, A2090T and C1944T in 23SrRNA were detected in two isolates with MIC=32μg/mL and 8μg/mL for LZD, respectively. In addition, 21 out of 25 clinical *M.fortuitum* isolates simultaneously showed following nine non-synonymous mutations in the coding protein of *rplD* for the both LZD susceptible and resistant isolates: Ala146Gly(GCG→GGC), Thr147Ser(ACC→AGC), Val156Ile(GTG→ATC), Ala161Thr(GCG → ACC), Lys167Arg(AAG → CGC), Ser207Ala(TCC → GCG), Glu212Gly(GAG→GGA), Val213Ala(GTG→GCG), Ala215Val(GCC→GTC) (Table 4).

**Table 4.** The MICs of LZD and *rplC*, *rplD* and *23srRNA* mutations against *M. fortuitum* isolates

| MIC of LZD (μg/ml) | Species (NO.) | RplC | RplD | 23SrRNA |
| --- | --- | --- | --- | --- |
| 4 | 1 | - | Ala146Gly+ Thr147Ser +<br>Val156Ile+ Ala161Thr+<br>Lys167Arg+ Ser207Ala+<br>Glu212Gly+ Val213Ala+<br>Ala215Val (1) |  |
| 8 | 2 |  | Ala146Gly+ Thr147Ser +<br>Val156Ile+ Ala161Thr+<br>Lys167Arg+ Ser207Ala+<br>Glu212Gly+ Val213Ala+<br>Ala215Val (1)<br>WT | C1944T(1) |
| 16 | 9 |  | Ala146Gly+ Thr147Ser +<br>Val156Ile+ Ala161Thr+<br>Lys167Arg+ Ser207Ala+<br>Glu212Gly+ Val213Ala+<br>Ala215Val (8) |  |
| 32 | 7 |  | Ala161Thr+ Lys167Arg+<br>Ser207Ala+ Glu212Gly+<br>Val213Ala+ Ala215Val(1)<br>Ala146Gly+ Thr147Ser +<br>Val156Ile+ Ala161Thr+<br>Lys167Arg+ Ser207Ala+<br>Glu212Gly+ Val213Ala+<br>Ala215Val (5) | A2090T(1) |
| >32 | 6 |  | Ala146Gly+ Thr147Ser +<br>Val156Ile+ Ala161Thr+<br>Lys167Arg+ Ser207Ala+<br>Glu212Gly+ Val213Ala+<br>Ala215Val (6) |  |

### Structural mapping of clinical mutants

For *M.massiliense* isolates, Ala177Pro in RplD was detected in 12 isolates, both in LZD susceptible and resistant isolates with MIC≥2μg/mL. To gain an insight into the functional relevance of RplC and RplD mutation, multiple sequences alignment of RplC and RplD homologues from different mycobacterial species were performed (Figure S1 and S2). The protein sequence of RplC and RplD in different mycobacterial species are highly conserved. In addition, we used *M. tuberculosis* RplD structure as a model to map *M.massiliense* RplD mutation(PDB ID:5V7Q) (Figure 5B). The structure shows that Ala177 is located in a high variable region between β3 and η2 and is far from the LZD binding site which indicates that this mutation may not be related to LZD resistance(Figure 3B). Next, we mapped the 23SrRNA functional mutations of *M.abscessus*, *M.massiliense* and *M.fortutium.* The results showed that except A2271 in *M.massiliense*, the other mutations including G2582, A2625 and T2650 were far from the catalytic center (Figure 3C).

**Figure 3.**
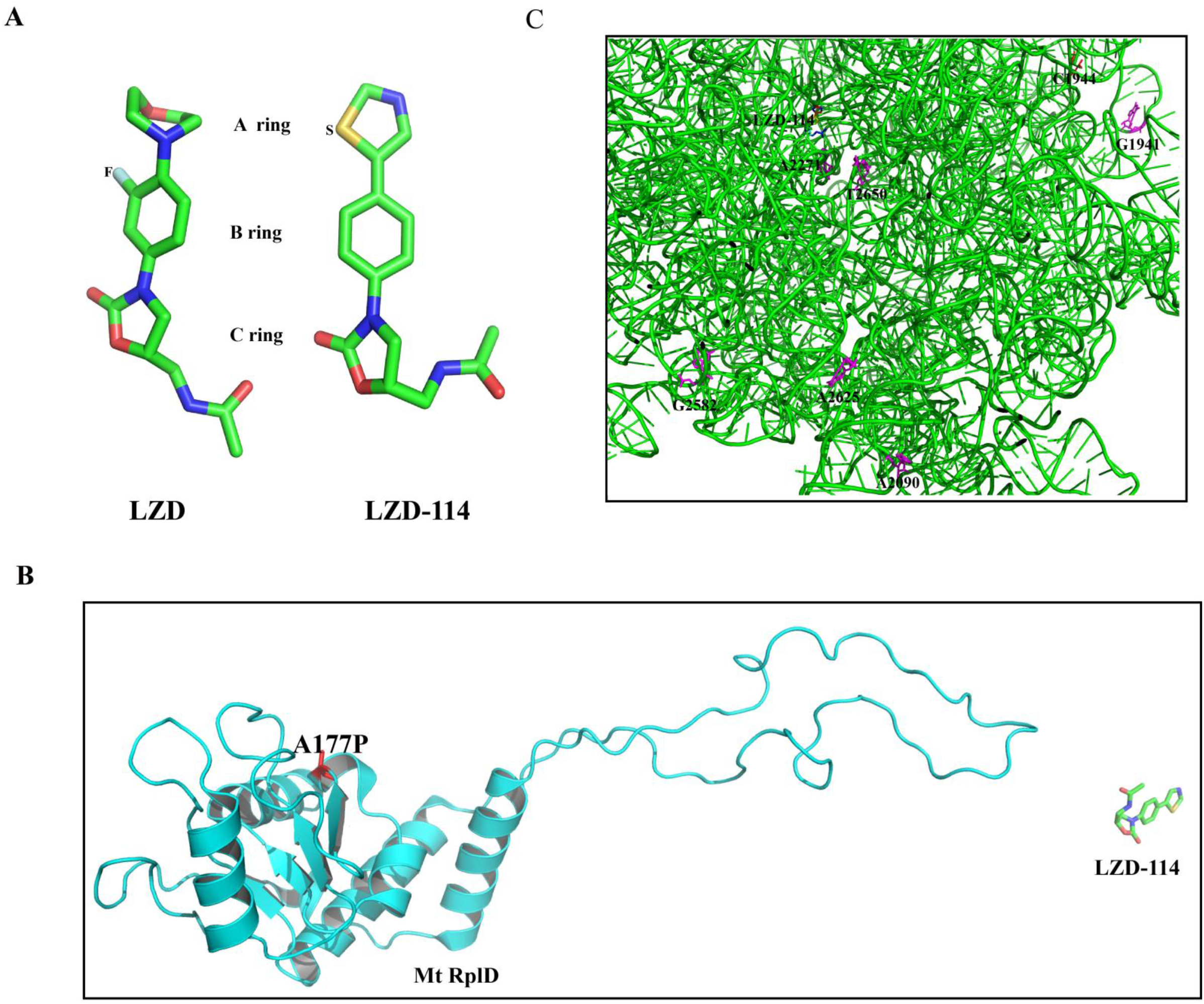
The structure of the ribosomal *23SrRNA* and *rplD*. (A)The structure of LZD and its analog LZD-114. (B)The structure of *rplD* and Ala177Pro mutations detected in *M.massiliense* isolated highlighted in red.(C) The structure of ribosomal *23SrRNA* and mutations detected in tested RGM clinical isolates were highlighted in red.

## DISCUSSION

The treatment of RGM infection is often very difficult because of their higher drug resistance rate than SGM and unavailability of highly potent drugs against them *in vitro*. *M.abscessus* complex and *M.fortuitum* are two most prevalent RGM species around the world. Infections due to *M.abscessus* carry a poor prognosis since this RGM is, for all the correct reasons, considered an “antibiotic nightmare”(17). Thus, identifying drugs that could work potently against *M.abscessus* is a priority. *M.massiliense* is a species that originally split from *M.abscessus* but they are located closely in the phylogenetic tree(18). The treatment response rates to clarithromycin-based antibiotic therapy are much higher in patients with *M.massiliense* than patients with *M.abscessus* lung disease(19). *M.fortuitum* is the main RGM responsible for extra-pulmonary disease, especially in cutaneous and plastic surgery-related infections(20). In contrast to *M.abscessus, M.fortuitum* infection has better prognosis due to some available effective drugs(21). However, its emerging drug resistance highlights the need for new and effective drugs(21–23). Several studies have verified the efficacy of LZD in MDR-TB or even in XDR-TB treatment (9, 13, 24). A few studies also proved its antibacterial activity against NTM species either *in vitro* or *in vivo*(25, 26). As a novel oxazolidinone prodrug, TZD exhibited greater potency than LZD against *M. tuberculosis*(6, 27) as well as against NTM(28, 29). Limited studies or no study has been performed to evaluate the efficacy of SZD and DZD against NTM species(28), whereas only a few studies provided preliminarily assessment of their potential usage in TB(14, 30, 31). In this study, we evaluated the efficacies of four oxazolidinones against the reference and clinical isolates of RGM to gain insights on their potential use for specific RGM species.

As new drugs, well recognized susceptibility testing methods for TZD, SZD and DTD have not been developed and the breakpoints to define drug resistance for them have never been discussed yet. Therefore, the MIC data of different RGM species against oxazolidinones still remain scarce. In this study, the four oxazolidinones exhibited promising activities *in vitro* against the recruited RGM reference stains. The absolute majority of species had MICs below 8μg/mL against the four drugs. However, different species presented non-uniform susceptibility patterns. The MIC distributions of *M.massiliense* had similar tendency to the *M.abscessus*, but the MICs of TZD were obviously higher than *M.abscessus*. In comparison with other oxazolidinones, the MIC values for TZD were the lowest for both *M.abscessus* and *M.massiliense.* Previous studies, including 170 isolates of RGM, showed equivalent or lower (1 to 8 fold) MIC_50_ and MIC_90_ values for TZD in contrast with LZD(29). Furthermore, TZD harbors several advantages over LZD in terms of tolerability, safety, dosing frequency, and treatment duration(32). Only a few studies have reported the clinical use of TZD for the treatment of NTM infections. Our results indicated that its usage seems reasonable for the treatment of infection caused by *M.abscessus* and *M.massiliense*. Among the 25 tested *M.fortuitum* isolates in our study, 22(22/25) strains had MICs of LZD at≥16 mg/L. Based on the CLSI criteria, these strains could be categorized as intermediate resistant or resistant strains, 52% (13/25) of them belong to resistant strains. Using the cutoff value of LZD as the tentative breakpoints, the susceptibility rate of *M.fortuitum* against TZD, SZD and DZD were 20%(5/25), 12%(3/25), 76%(19/25), respectively. DZD exhibited the best antimicrobial activity against the *M.fortuitum*. However, whether this *in vitro* outcome reflects the *in vivo* efficacy or not, requires further investigation.

A major limitation of this study was that no recommended breakpoint of different NTM species against TZD, SZD or DZD had been proposed previously. Beside *in vitro* MIC distributions, the breakpoint determination also correlates with clinical treatment response and pharmacokinetic/pharmacodynamics (PK/PD) data including drug dose. The clinical trial on these new oxazolidinones are either unavailable or very limited. A few studies have been performed on the pharmacokinetic analysis of these drugs. Generally, all the drugs were well-tolerated, and the C_max_ were highly dose-dependent. Recently, Choi et al demonstrated that, after multiple doses of TZD up to 1200mg twice daily for 21days, the peak serum concentration was 16.3μg/mL, which is comparable with peak serum concentrations of LZD=12.5μg/mL at the dosage of 300mg twice daily(15, 33). In another study, a single 800mg dose of DZD under fasting condition acquired C_max_ at 11.74μg/mL(34). STD presented superior efficacy than LZD against experimental murine model of tuberculosis. The C_max_ of its major metabolite PNU-101603, which contributes to its activity, was 6.46μg/mL at given dose of 1200mg QD (40). However, since the optimal dosage of these next-generation oxazolidinones is still under investigation, the appropriate breakpoints for the susceptibility definition of these drugs remain beyond known.

LZD works by binding to the peptidyl transferase center of the 50S ribosomal subunit, which is composed of 5S and 23S rRNAs and 36 riboproteins (L1 through L36)(35). Recently, the Cryo-EM structure of the large ribosomal subunit from *M.tuberculosis* bound with a potent LZD analog (LZD-114) was determined(36). LZD-114 is similar with LZD in C ring but different in A and B ring that lacks a fluorine group in the B-ring while the original morpholine ring is replaced by a thiazole ring in the A-ring(Figure 3A). The LZD-114 also was bound in the same pocket and in a similar orientation to LZD in other species(37, 38). The structure showed that *rplC* encoded ribosomal protein L3 and *rplD* encoded ribosomal protein L4 bind directly to 23S ribosomal RNA and was placed relatively close to the LZD binding site on the ribosomes, suggesting that the mechanism for reduced susceptibility may include structural perturbation of the LZD binding site (PDB:5V7Q). Furthermore, previous studies demonstrated that mutations in *rplC* and *rplD* could lead to LZD resistance in *M.tuberculosis*(12, 39). However, there is no non-synonymous mutation in *rplC* against the tested RGM. Ala77Pro mutation was detected in *rplD* which is located in variable site and is far away from LZD-binding site, as shown by the sequence alignment. Except A2271G mutation in 23SrRNA in *M.massiliense* that was closer to binding site of LZD, other mutations are far from the LZD-binding site. Our results combining MIC test, gene mutation and structure based analysis showed there is no obvious correlation between riboproteins mutations (*rplC* and *rplD*) and LZD resistance identified in this study in the RGM species. Mutations located in the LZD binding site may cause LZD resistant. Hence, *rplC*, *rplD* and *23srRNA* homologues might not be the only target for LZD to explore its bacteriostatic activity.

In conclusion, this study demonstrated that oxazolidinones have good *in vitro* activities against the overwhelming majority of RGM species. The efficacies of the four oxazolidinones were variable against different species. TZD showed strongest antimicrobial activity against *M.abscessus* and *M.massiliense*, while DZD owned the strongest activity against *M.fortuitum*. The data provided important insights on the possible clinical applications of oxazolidinones to treat RGM infections.

## MATERIAL AND METHODS

### Ethics statement

As the study only concerned laboratory testing of mycobacteria without the direct involvement of human subjects, ethics approval was not sought.

### Reference strains and clinical isolates

The mycobacterial reference strains stored in the Bio-bank in Beijing Chest Hospital (Beijing, China) were tested against LZD, TZD, SZD and DZD *in vitro*, including 32 RGM species. These reference strains were obtained either from the American Type Culture Collection (ATCC) or from the German Collection of Microorganisms (DSM). The species constitution of these reference strains are listed in Table 1. *M.massiliense* reference strain was not included due to its absence in our stock. One-hundred fifteen isolates of RGM were recruited in Beijing chest hospital from 2016 to 2018 that included 49 *M.abscessus*, 35 *M.massiliense*, 25 *M.fortuitum*. The species constitution of the remaining 6 isolates is presented in Table 2.

All of the 115 RGM clinical strains were isolated from tuberculosis suspected patients. The strains were classified as RGM preliminarily with p-nitrobenzoic acid containing medium, and then were identified by gene sequencing as indicated for each species by *16S rRNA*, *hsp65, rpoB*, *16-23S rRNA* internal transcribed spacer sequencing (40). All the isolates were stored at −80°C and sub-cultured on LJ medium before performing drug susceptibility test.

### Minimal inhibitory concentration (MIC) testing

TZD phosphate and LZD were purchased from Toronto Research Chemicals and Sigma-aldrich, respectively. SZD and DZD were purchased from Shanghai Biochempartner Co., Ltd (Shanghai, China) and JHK BioPharma, respectively. Oxazolidinones were dissolved in dimethyl sulfoxide (DMSO). Stock solutions were aseptically prepared at concentrations of 2.56 mg/mL. Broth microdilution method was performed according to the guidelines of Clinical and Laboratory Standards Institute (CLSI)(41). Cation-adjusted Mueller-Hinton broth (CAMHB) was used for MIC test. The inoculum was prepared with fresh culture grown on Lowenstein-Jensen medium. The broth microdilution format was set up as 2-fold dilution, the concentrations of all the tested drugs ranged from 0.063μg/mL to 32μg/mL. Briefly, a bacterial suspension of 0.5 McFarland standard was prepared, and then diluted and inoculated into 96-well microtiter plate to achieve final bacterial load at 10^5^ colony forming unit (CFU) per well. Plates were then incubated at 37°C for 3 days for RGM. 70μL solution containing 20μL AlamarBlue (Bio-rad) and 50μL Tween80 (5%) was added to each well and incubated for 24 h at 37 °C before assessing color development. A change from blue to pink or purple indicated bacterial growth (42). The MIC was defined as the lowest concentration of antibiotic that prevented a color change from blue to pink.

The breakpoint of LZD was adopted from the CLSI document M24-A2 (susceptible:≤8 mg/L; intermediate resistant: 16 mg/L; resistant:≥32 mg/L) (16). Since no well-recognized breakpoint has been proposed for TZD, SZD or DZD, a preliminary data analysis was performed for them referring the breakpoint of LZD.

### Mutations conferring oxazolidinones resistance and protein Alignment

Sequencing of PCR products was performed using the Sanger method with primers designed to be specific for *rplC*, *rplD* and *23S rRNA*. We used previously described primers for *23S rRNA*(43), and design new primers for *rplC, rplD* sequencing. The primers used in this study are listed in Table S1 in the supplemental material and were synthesized by Tsingke Biotech Co. (Beijing, China). The *rplC* and *rplD* gene of the reference strains of three RGM species plus *M.tuberculosis* were also sequenced, mutation was defined in contrast with the sequences of the reference strains. The sequences of *M.massiliense* were adopted from website for alignment. The amplification products were sequenced by Tsingke Company (Beijing, China). Multiple sequence alignment of the homologous proteins was performed using the Clustal Omega software. Structure-based multiple sequence alignment was performed with ESPript 3 based on the crystal structure of RplC and RplD protein of *M.tuberculosis* from the following URL:http://espript.ibcp.fr/ESPript/ESPript/.

#### Quality control

The MIC for quality control strains was determined using each lot of the prepared microtiter plates, and the results for LZD were within the expected range.

## ACKNOWLEDGEMENT

This study was supported by research funding from the Infectious Diseases Special Project, Ministry of Health of China (2018ZX10302302-004-005, 2018ZX10201301302-004.) the Natural Science Fund of China (81672065, 81802057), Beijing Municipal Administration of Hospitals Clinical Medicine Development of Special Funding Support (ZYLX201824) and Beijing Municipal Administration of Hospitals’ Ascent Plan (DFL20181602).

## TRANSPARENCY DECLARATIONS

None to declare.

---

**Table S1. Primer sets used for target genes in this study**

**FigureS1. Sequence alignment *rplC* homologue proteins.** Alignment of the amino acid sequences of *M. tuberculosis*, *M.abscessus, M.massiliense, M.chelonae, M. fortuitum and M.smegmatis.* The topology of the *rplC* encoded protein of *M. tuberculosis* is shown at the top. Red boxes with white letters indicate a single, fully conserved residue. Blue frames indicate highly conserved residues. *β*-Strands are rendered as arrows.

**FigureS2. Sequence alignment *rplD* homologue proteins.** Alignment of the amino acid sequences of *M. tuberculosis*, *M.abscessus, M. massiliense, M.chelonae, M. fortuitum and M.smegmatis.* The topology of the *rplD* encoded protein of *M. tuberculosis* is shown at the top. Red boxes with white letters indicate a single, fully conserved residue. Blue frames indicate highly conserved residues. *β*-Strands are rendered as arrows.

